# Analysis of *Drosophila* cardiac hypertrophy by micro-computerized tomography for genetic dissection of heart growth mechanisms

**DOI:** 10.1101/2021.01.22.427777

**Authors:** Courtney E Petersen, Benjamin A Tripoli, Todd A Schoborg, Jeremy T Smyth

## Abstract

Heart failure is often preceded by pathological cardiac hypertrophy, a thickening of the heart musculature driven by complex gene regulatory and signaling processes. The *Drosophila* heart has great potential as a genetic model for deciphering the underlying mechanisms of cardiac hypertrophy. However, current methods for evaluating hypertrophy of the *Drosophila* heart are laborious and difficult to carry out reproducibly. Here we demonstrate that micro-computerized tomography (microCT) is an accessible, highly reproducible method for non-destructive, quantitative analysis of *Drosophila* heart morphology and size. To validate our microCT approach for analyzing *Drosophila* cardiac hypertrophy, we show that expression of constitutively active Ras (*Ras85D*^*V12*^), previously shown to cause hypertrophy of the fly heart, results in significant thickening of both adult and larval heart walls when measured from microCT images. We then show using microCT analysis that genetic upregulation of store-operated Ca^2+^ entry (SOCE) driven by expression of constitutively active Stim (Stim^CA^) or Orai (Orai^CA^) proteins also results in significant hypertrophy of the *Drosophila* heart, through a process that specifically depends on Orai Ca^2+^ influx channels. Intravital imaging of heart contractility revealed significantly reduced end diastolic dimensions in Stim^CA^ and Orai^CA^ expressing hearts, consistent with the hypertrophic phenotype. These results demonstrate that increased SOCE activity is an important driver of hypertrophic cardiomyocyte growth, and demonstrate how microCT analysis combined with tractable genetic tools in *Drosophila* can be used to delineate molecular signaling processes that underlie cardiac hypertrophy and heart failure.

**NEW AND NOTEWORTHY:** Genetic analysis of cardiac hypertrophy in *Drosophila* holds immense potential for the discovery of new therapeutic targets to prevent and treat heart failure. However, this potential has been hindered by a lack of rapid and effective methods for analysis of heart size in flies. Here we demonstrate that analysis of the *Drosophila* heart with micro-computerized tomography yields accurate and highly reproducible heart size measurements that can be used to efficiently analyze heart growth and cardiac hypertrophy in *Drosophila*.

## INTRODUCTION

Pathological cardiac hypertrophy occurs when the heart enlarges due to chronic functional overload. While initially compensatory, unmitigated hypertrophy eventually transitions to heart failure^1-4^. There are no cures for pathological cardiac hypertrophy or heart failure, and better understanding of the cellular and molecular mechanisms that contribute to maladaptive cardiomyocyte growth is therefore essential.

Cardiac hypertrophy is driven by complex integration of multiple signaling pathways including the Ca^2+^-calcineurin-NFAT, Ras-Raf-MAPK, and PI3K signaling axes^2^. Resolution of this complexity will benefit from molecular pathway dissection in a genetically tractable model like *Drosophila melanogaster*. The *Drosophila* heart is a linear tube of cardiomyocytes that runs along the dorsal midline of the adult abdomen, and it pumps hemolymph throughout an open circulatory system^5-7^. A major advantage of using *Drosophila* for cardiac research is the ability to rapidly engineer multiple genetic alterations in the same animal for integrated signaling pathway analysis. These approaches have shown that both Ras-Raf-MAPK and calcineurin-regulated signaling pathways drive cardiac hypertrophy in *Drosophila*^8-10^, demonstrating that hypertrophic signaling in flies is well conserved with mammals. However, the full potential of *Drosophila* for cardiac hypertrophy research has been limited by a lack of readily accessible and reproducible methods for analyzing hypertrophic phenotypes in flies.

Key phenotypic indicators of cardiac hypertrophy include increased heart wall thickness and reduced luminal diastolic and systolic dimensions. Methods to analyze *Drosophila* heart contractile parameters including end-diastolic and systolic dimensions are well established^11^. However, analyses of *Drosophila* heart wall thickness are limited and have relied on measurements derived from histopathological approaches involving physical sectioning of fixed and embedded flies^9, 10, 12, 13^. Histopathology has several drawbacks that limit throughput, reproducibility, and accuracy. First, it is very time-consuming and laborious, and this limits the feasibility of processing large numbers of samples required for genetic pathway analyses. Second, achieving reproducible sectioning through the same region of the heart and in the same orientation is challenging. And third, physical sectioning through the heart yields relatively thick sections of 8-10 μm, which can limit the accuracy of individual measurements.

Several recent studies have described micro-computerized tomography (microCT) for quantitative analysis of internal organs in *Drosophila*^14-19^, and our study aimed to determine the feasibility of microCT for quantitative analysis of cardiac hypertrophy in flies. MicroCT is an x-ray imaging methodology whereby the sample is incrementally rotated to generate a series of projection images that encompasses the full sample volume. Computerized algorithms then generate three-dimensional (3D) tomogram reconstructions of the sample with isotropic resolution. Due to the small size of *Drosophila*, an entire animal at any developmental stage can be imaged in 3D by microCT in a single acquisition sequence, allowing visualization of internal organs in any orientation and location with exceptional clarity and resolution^19^. Importantly, microCT measurements can be reproducibly made from the same location within a specific organ or structure without imprecise physical sectioning. MicroCT also involves straightforward fixation and staining protocols, and multiple animals can be processed at a time. Reproducible measurements of wildtype *Drosophila* hearts have been made using microCT^19^; however, whether microCT can be used to analyze pathological changes to heart architecture such as hypertrophy has not been determined. Integration of microCT with combinatorial *Drosophila* genetic tools has immense potential to provide new insights into molecular mechanisms of pathological heart remodeling.

Here we demonstrate that analytical microCT approaches accurately and reproducibly reports hypertrophy of the *Drosophila* heart. As proof-of-principle, we show using microCT that expression of constitutively-active Ras, previously demonstrated to cause cardiac hypertrophy in *Drosophila* by histopathological analyses^8^, results in significantly increased heart wall thickness in adult and larval flies. We then demonstrate using genetic approaches, microCT measurements, and contractility analysis that upregulation of store-operated Ca^2+^ entry (SOCE) drives cardiac hypertrophy in *Drosophila*. SOCE is a highly conserved Ca^2+^ signaling mechanism that couples influx of extracellular Ca^2+^ to depletion of endo/sarcoplasmic reticulum (E/SR) Ca^2+^ stores. SOCE is mediated by Ca^2+^-sensing Stim proteins in the E/SR and Orai Ca^2+^ influx channels in the plasma membrane^20-22^. SOCE upregulation in mammalian cardiomyocytes is necessary and sufficient for pressure-overload induced cardiac hypertrophy, likely due to SOCE activation of calcineurin signaling^23-25^. However, mechanisms that regulate SOCE in cardiomyocytes, and how SOCE integrates with other hypertrophic signaling processes are poorly understood. Our results establish a new model of SOCE-driven cardiac hypertrophy that combines the power of *Drosophila* genetics with robust analysis of fly heart wall thickness with microCT. This microCT approach is straightforward, accessible, and readily adaptable to the renowned genetic tools in *Drosophila*, creating a powerful experimental platform for delineating the complex cellular and genetic processes that drive pathological cardiac hypertrophy.

## MATERIALS AND METHODS

### Fly Stocks

*w1118* (3605), *UAS-Ras85D*^*V12*^ (64195), and *Orai* RNAi (53333) were from the Bloomington *Drosophila* Stock Center. *tinC-GAL4* was from Dr. Manfred Frausch (Friedrich Alexander University). Flies expressing tdTomato under control of the cardiomyocyte-specific *R94C02* enhancer^26^ (*CM-tdTom*) were from Dr. Rolf Bodmer (Sanford Burnham Prebys Institute). *CM-tdTom*^*2*^ and *CM-tdTom*^*3*^ are second and third chromosome insertions, respectively. Wildtype *Orai* (*Orai*^*WT*^) and constitutively-active *Orai* (*Orai*^*CA*^) were from Dr. Gaiti Hasan (National Center for Biological Sciences, Bangalore, India). Flies that express wildtype *Drosophila Stim* (*Stim*^*WT*^) under *UASp* control were generated by cloning the cDNA sequence of *Drosophila Stim* Isoform A from plasmid LD45776 (*Drosophila* Genomics Resource Center). *Stim* cDNA was inserted into vector pPWG (Carnegie *Drosophila* Gateway Vector Collection), which introduces a C-terminal EGFP tag and upstream *UASp*. For constitutively-active *Stim* (*Stim*^*CA*^), site-directed mutagenesis (Stratagene QuikChange XL) was used to change aspartic acids 155 and 157 to alanines using *StimWT* as template. Sequence confirmed plasmids were sent to BestGene for embryo injection.

### Micro-computerized Tomography (microCT)

The following microCT methods were adapted from Schoborg et al^19, 27^:

#### Adult Labeling

5-20 adult flies were anesthetized with CO2 and transferred to a 1.5 ml Eppendorf tube containing 1 ml of phosphate buffered saline + 0.5% Triton-X 100 (0.5% PBST). Tubes were gently inverted, then incubated 5 minutes at room temperature (RT) to remove wax cuticles. Flies were transferred to tubes containing 1 ml Bouin’s fixative (5% acetic acid, 9% formaldehyde, 0.9% picric acid; Sigma) for 24 hours. Samples were washed 3 × 30 minutes on a shaker in 1 ml microCT Wash Buffer (0.1 M Na2HPO4/NaH2P04 + 1.8% Sucrose, pH 7.0), followed by staining with 1 ml of 0.1 N iodine-iodide solution (Lugol’s solution) for 48 hours. Flies were then washed twice with ultrapure water and stored at RT for up to one month.

#### Larval Labeling

Third-instar larvae were placed in 1.5 ml Eppendorf tube with 1 ml 0.5% PBST. Tubes were heated to 100 ° C for 20 seconds, then cooled to RT for 5 minutes. Larvae were then fixed in Bouin’s solution for 24 hours and washed as for adults. Larval cuticles were then punctured at anterior and posterior ends with a microdissection needle to allow penetration of the labeling solution. Punctured larvae were incubated in Lugol’s solution for 48 hours, washed twice with ultrapure water, and stored at RT for up to one month.

#### Sample Mounting and Scanning

Individual adults and larvae were placed head-down in heat sealed 10 µl micropipette tips containing ultrapure water, and a dulled 20-guage needle was used to gently lower the animals until they fit snuggly in the taper of the pipette tip. Parafilm was wrapped around the base of the micropipette tip to prevent leakage. Samples were secured to the stage of the microCT scanner, pipette base down, using mounting putty. Samples were scanned with a Bruker SkyScan 1172 scanner controlled by Skyscan software operated on a Dell computer. The following X-ray source voltage and current settings were used: 40 kV, 110 μA, and 4 W. A Hamamatsu 10 Mp camera with 11.54 μm pixels coupled to a scintillator was used to collect X-rays and convert to photons. Fast scans were at a camera resolution of 2.85 μm, and slow scans at 1.15 μm with 360 degrees of sample rotation. Frame averaging was four for all scans. At the scanner and reconstruction settings used, the SkyScan 1172 produces tomograms with in-plane and axial spatial resolutions of 5-9 μm as determined by the QRM-MicroCT-BarPattern NANO (QRM GmbH, Möhrendorf Germany).

#### Reconstruction

Tomograms were generated using NRecon software (Bruker, v1.7.0.4). The built-in shift correction function of NRecon, which uses reference scans to compensate for sample movement during scanning, was used for image alignment and reconstruction. Remaining misalignment was manually fine-tuned using the misalignment compensation function. Ring artifact correction was set to max (50) and beam hardening was 0%.

#### Heart Wall Thickness Measurements

Reconstructed microCT tomogram series were imported into FIJI (ImageJ v1.53c, NIH) and viewed using the Orthogonal Views function to locate hearts in XY, XZ, and YZ orientations. Using images that showed the largest opening of the conical chamber in cross-section, the thickest heart wall sections were measured with line segments in FIJI. An average of ten measurements from five slices was used to represent each animal. All measurements were made blinded to genotype.

### Contractility Analysis

Intravital fluorescence imaging of hearts expressing CM-tdTom was carried out as previously described^28^. Seven day-old adult females were briefly anesthetized with CO2 and adhered dorsal side down to glass coverslips with Norland Optical Adhesive cured with a 48-watt UV LED source (LKE) for 60 seconds. Animals recovered for 10 minutes before imaging. Hearts were imaged through the dorsal cuticle at 200 frames-per-second for twenty seconds using an ORCA-Flash4.0 V3 sCMOS camera (Hamamatsu) on a Nikon-Ti2 inverted microscope controlled with Nikon Elements software. 550 nm excitation light was from a Spectra-X illuminator (Lumencor) and emission was collected through a 555-635 nm band-pass filter. To generate M-modes, a 1-pixel wide line was drawn through the heart in the A2 segment, and fluorescence intensity along this line was plotted using Multi-Kymograph in FIJI. End-diastolic dimensions (EDD) and end-systolic dimensions (ESD) were calculated from the M-modes by measuring the distance between the heart walls at full relaxation and contraction, respectively. An average of five EDD and ESD measurements was calculated from each trace. Heart rate was calculated by counting the number of systoles over 20 seconds. FS was calculated as ((EDD-ESD) / EDD) x 100. All measurements were made blinded to genotype.

### Statistical Analyses

Plots and statistical analyses were done with GraphPad Prism. Contractility and heart wall measurements were analyzed by unpaired t-test or one-way ANOVA with Tukey’s Multiple Comparisons Test. Statistical significance was at p < 0.05.

## RESULTS

### Quantitative microCT analysis of cardiac hypertrophy in adult *Drosophila*

We began validating our microCT methodology by analyzing iodine-labelled hearts in *w1118* control adult flies. We imaged adults by microCT at the highest resolution settings for which the entire animal fits within the detector’s field of view. These “slow-scans” generate tomograms with 1.15 µm^2^ pixels and an actual spatial resolution of 5-6 μm (See Methods). Figure 1A shows a 3D reconstruction of a wildtype *w1118* adult female fly generated from slow-scan microCT tomograms. The heart tube runs along the dorsal abdominal wall midline and is most easily identified in 2D at the anterior and dorsal-most region of the abdomen (Figure 1B). This anterior-most segment of the heart, which exhibits the largest luminal dimensions of the heart tube, is known as the conical chamber (Figure 1B-D). It can be identified in the XY view by locating the prominent dorsal longitudinal muscles (DLMs) of the thorax (Figure 1C) and following them posteriorly to where they narrow and the abdomen begins. Because of the large luminal space of the conical chamber and the exceptional contrast and clarity provided by microCT, XY cross-sections through the conical chamber are ideal for visualizing and measuring the heart walls (Figure 1D).

**Figure 1.**
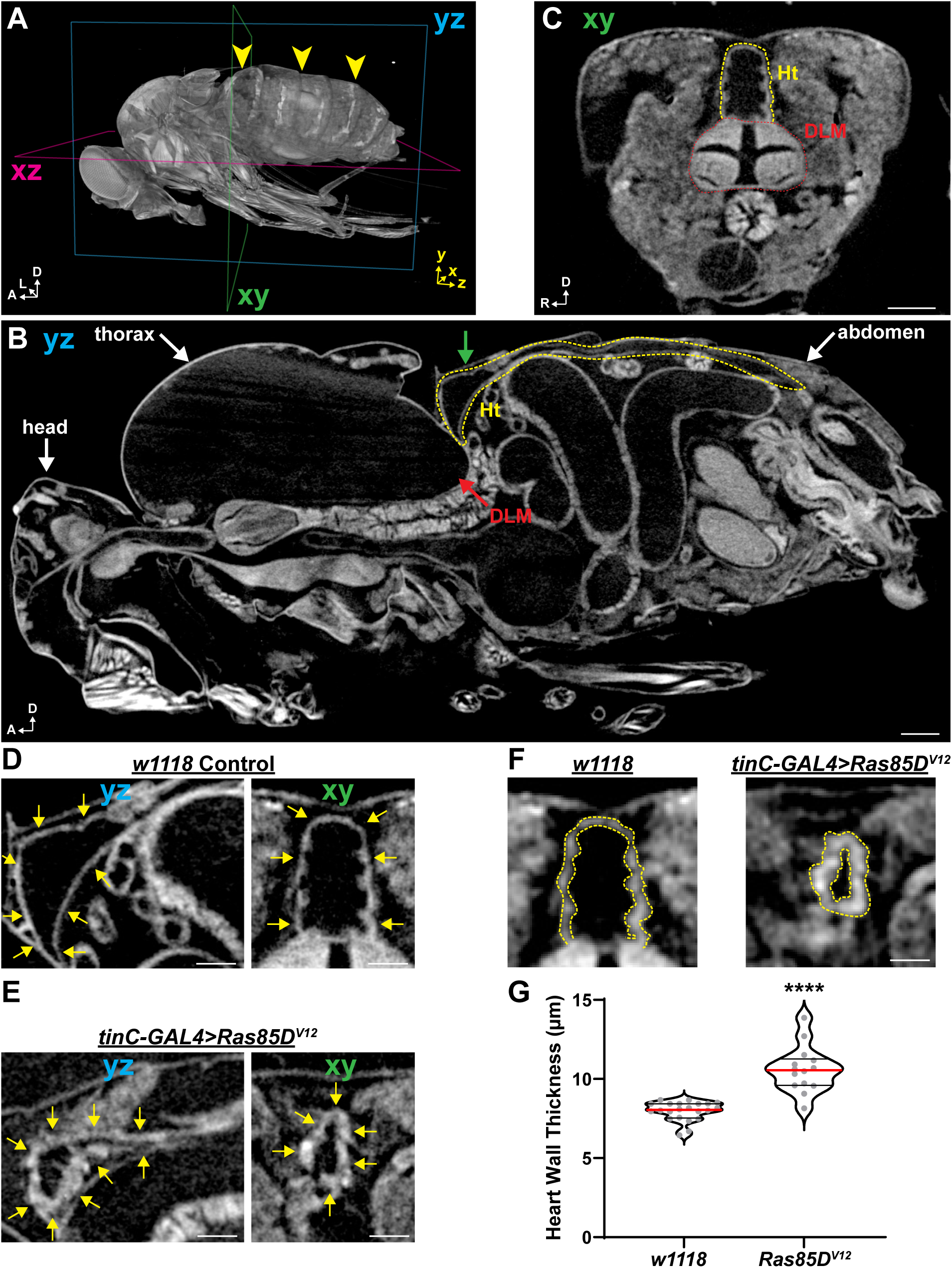
MicroCT analysis of cardiac hypertrophy in adult *Drosophila*. **A**. 3D reconstruction of a seven day-old female fly generated from slow-scan microCT tomograms. XY, XZ, and YZ imaging planes are denoted by colored rectangles, and anatomical anterior (A), left (L), and dorsal (D) are indicated by white arrow coordinates. Yellow arrowheads indicate the location of the heart along the midline of the dorsal abdomen. **B**. Slow-scan YZ-microCT tomogram at the midline of a seven day-old *w1118* female fly, providing a longitudinal view of the animal. Head, thorax, and abdomen are indicated for orientation. The yellow dashed line outlines the heart (Ht) longitudinally from anterior to posterior of the dorsal abdomen. The thoracic dorsal longitudinal muscles (DLM) are also indicated; note the region near the red arrow where the posterior DLMs are just beneath the anterior start of the heart, as this overlap is useful for identifying the heart conical chamber. **C**. Single XY-tomogram from the location indicated by the green arrow in panel B, showing the anterior abdomen in cross-section. The heart conical chamber in cross-section (outlined in yellow dashes) is easily identified just dorsal to the small bundle of DLMs (outlined by red dashes). **D**. Enlargements from panels B and C showing the heart conical chamber in longitudinal (yz; left) and cross-section (XY; right) orientations. Yellow arrows denote the heart walls. **E**. Longitudinal (YZ; left) and cross-section (XY; right) images from slow-scan microCT tomograms of a *tinC-GAL4*>*Ras85D*^*v12*^ seven day-old female fly. Yellow arrows denote the heart walls. Note the significant thickening of the heart walls compared to the *w1118* control in panel D. **F**. Representative cross-section images of heart conical chambers from fast-scan microCT tomograms of the same w*1118* and *tinC-GAL4*>*Ras85D*^*v12*^ animals shown in panels B-E. Dashed yellow lines indicate the inner and outer boundaries of the heart walls. **G**. Plot of heart wall thickness measurements from fast-scan microCT tomograms of *w1118* control and *tinC-GAL4*>*Ras85D*^*v12*^ seven day-old females. Each circle represents a single animal measurement. Red lines: median; black lines: quartiles. ****p<0.0001, unpaired t-test. Scalebars: B-C, 100 µm; D-F, 50 µm.

We next determined whether microCT can be used to quantitatively analyze hypertrophy of the adult *Drosophila* heart by expressing a constitutively-active Ras mutant (*Ras85D*^*V12*^) previously shown to induce *Drosophila* cardiac hypertrophy^8^. Heart-specific expression of *Ras85D*^*V12*^ using *tinC-GAL4* resulted in strikingly apparent thickening of heart walls in longitudinal and cross-section slow-scan images of the conical chamber compared to *w1118* controls (Figure 1E, Supplemental Video 1: https://figshare.com/s/d480f3f7ab20165d7f04, Supplemental Video 2: https://figshare.com/s/349e8616ca32ec94a9ea). These slow-scans provide maximal resolution for whole-animal imaging; however, slow-scans require 6-8 hours to complete and are not practical for generating large datasets for quantitative analysis. To facilitate higher throughput for quantitative measurements, we next tested “fast-scans” that require 20-30 minutes per animal and generate tomograms with 2.85 µm^2^ pixels. Conical chamber cross-sections from fast-scans also clearly exhibited visual thickening of heart walls in *Ras85D*^*V12*^ compared to *w1118* control hearts (Figure 1F). Direct measurement of heart wall thickness at the conical chambers from fast-scan, cross-section images from 22 *w1118* control animals yielded a mean of 7.97 ± 0.13 μm (Figure 1G, ± SEM). Importantly, these measurements from microCT fast-scans were nearly identical to previously published measurements from histopathological preparations of control hearts^8-10^. Moreover, the low variance suggests a high degree of reproducibility of these microCT-based measurements. Measurements from 14 *Ras85D*^*V12*^-expressing hearts revealed a significant, 33% increase in heart wall thickness compared to *w1118* (Figure 1G). These results demonstrate that our microCT methodology accurately reports the significant increase in heart wall thickness caused by *Ras85D*^*V12*^ expression, suggesting that microCT imaging is a robust method for analyzing adult *Drosophila* cardiac hypertrophy.

### MicroCT analysis of larval heart wall thickness

The majority of cardiomyocyte growth in *Drosophila* occurs during the larval developmental stages, making the larval heart useful for studying both physiological and pathological heart growth mechanisms^7, 29^. As with adult flies however, few tools are currently available for quantitative analysis of larval heart growth parameters such as heart wall thickness. We therefore assessed microCT for evaluating heart wall thickness in larvae. The heart spans the posterior third of third-instar larvae and is situated just under the dorsal cuticle (Figure 2 A,B). In cross-section, the heart appears as a round, thin-walled structure at the dorsal mid-line (Figure 2C). It is important not to mistake the heart for the two trachea, which are also round tubes found near the dorsal mid-line. The best way to identify the trachea is to find the two spiracles at the posterior end of the animal (indicated in Figure 2B), and to then follow the spiracles into the animal where they become continuous with the trachea. The heart is then found between and just dorsal to the two trachea (best seen in Figure 2D). In fast-scans, *tinC-GAL4*-driven *Ras85D*^*V12*^ expression resulted in visible thickening of larval heart walls and narrowing of the heart lumen compared to *w1118* controls. (Figure 2D). Direct measurements of heart wall thickness in cross-section revealed a 26% increase in *Ras85D*^*V12*^ compared to *w1118* hearts, from 7.24 ± 0.10 μm to 9.15 ± 0.26 µm, respectively (mean ± SEM; Figure 2E). These results demonstrate that microCT generates images of *Drosophila* larvae with sufficient resolution and sensitivity to detect hypertrophy of the larval heart.

**Figure 2.**
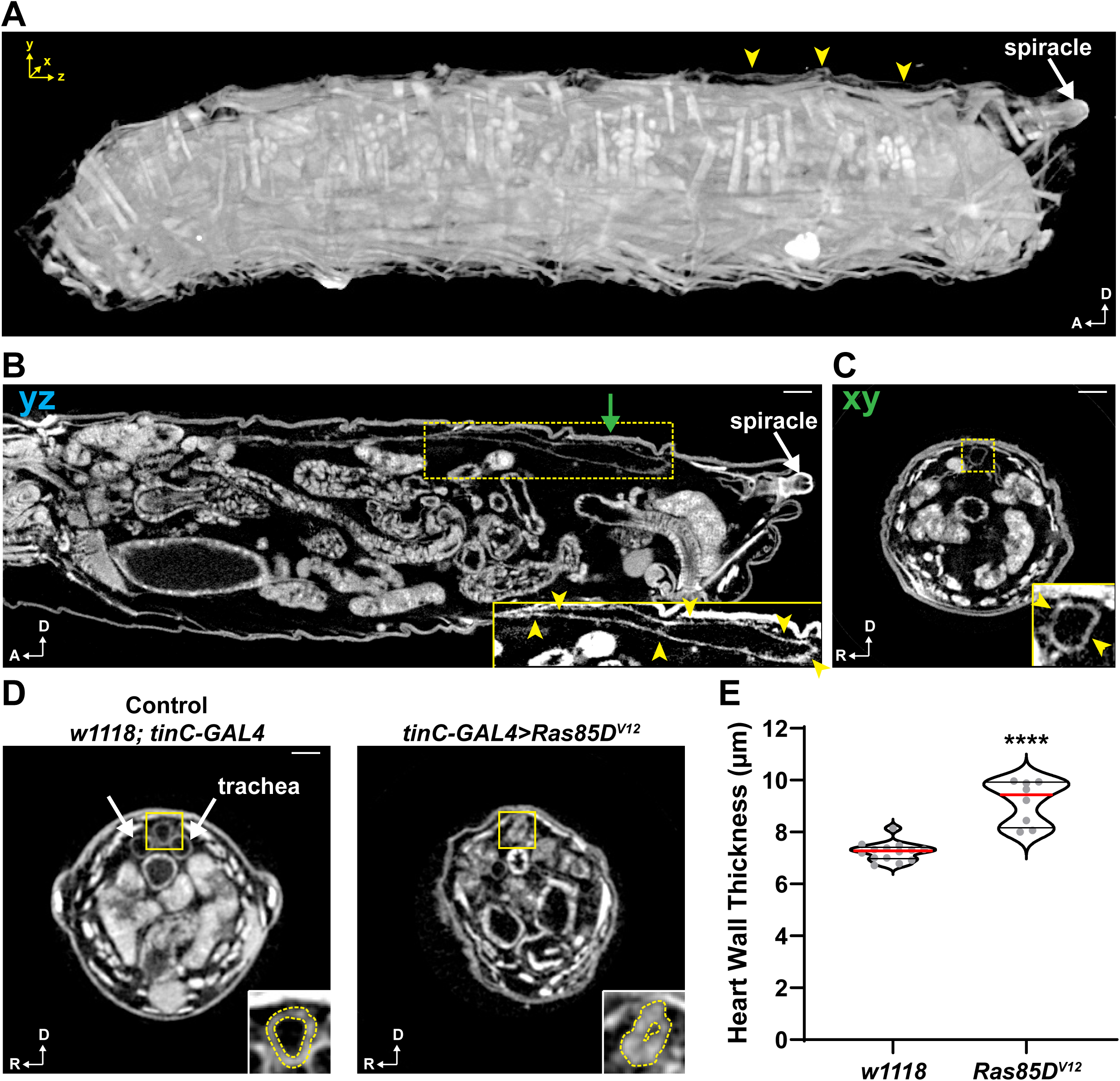
MicroCT analysis of cardiac hypertrophy in *Drosophila* third-instar larvae. **A**. 3D reconstruction of a third-instar larva generated from microCT tomograms. Anatomical anterior (A) and dorsal (D) are indicated by white arrow coordinates. Yellow arrowheads indicate the location of the heart along the dorsal midline in the posterior third of the animal. **B**. Single slow-scan YZ-microCT tomogram at the midline of a *w1118* third-instar larva, providing a longitudinal view of the animal’s internal organs. Note that the larval anterior was not captured because the larva was larger than the detector’s field of view. The yellow dashed rectangle denotes the heart seen longitudinally, and the region within the rectangle is enlarged and contrast-enhanced in the lower right inset. Yellow arrowheads in the inset point to the heart walls. **C**. Single XY-tomogram from the location indicated by the green arrow in panel B, showing the larva in cross-section. The yellow dashed square denotes the heart, and the region within the square is enlarged and contrast-enhanced in the lower right inset. Yellow arrowheads in the inset point to the heart walls. **D**. Representative cross-section images of control (*w1118*; *tinC-GAL4*) and *tinC-GAL4*>*Ras85D*^*v12*^ third-instar larvae. Yellow squares denote the hearts as well as the areas enlarged in the bottom right insets. Dashed yellow lines in the insets indicate inner and outer boundaries of the heart walls. **E**. Plot of heart wall thickness measurements from fast-scan microCT tomograms of control (*w1118*; *tinC-GAL4*) and *tinC-GAL4*>*Ras85D*^*v12*^ third-instar larvae. Each circle represents a single animal measurement. Red lines: median; black lines: quartiles. ****p<0.0001, unpaired t-test. Scalebars: 100 µm.

### SOCE upregulation results in cardiac hypertrophy

SOCE upregulation in cardiomyocytes is required for induction of pathological cardiac hypertrophy in vertebrates^23-25, 30, 31^. *Drosophila* may be an important model to elucidate how SOCE regulates cardiac hypertrophy, but whether SOCE upregulation results in hypertrophy of the *Drosophila* heart is not known. We therefore used microCT to evaluate whether SOCE upregulation due to expression of constitutively-active *Stim* and *Orai* mutants results in increased *Drosophila* heart wall thickness. Constitutively-active Stim (Stim^CA^) contains two aspartate to alanine substitutions within the EF-hand domain, causing Stim^CA^ to function in a Ca^2+^-unbound, fully-active state^32^. Constitutively-active Orai (Orai^CA^) has a glycine to methionine substitution in the channel hinge region, forcing the channel into an open conformation^33^. Heart specific *Stim*^*CA*^ and *Orai*^*CA*^ expression resulted in strikingly thicker heart walls in adults compared to controls when viewed in cross section (Figure 3A,B; Supplemental Video 3: https://figshare.com/s/f8769204768570064558, Supplemental Video 4: https://figshare.com/s/ebb84e37676ca8a1f22d), similar to heart wall thickening seen with *Ras85D*^*V12*^ (compare to Figure 1F). The increase in heart wall thickness over controls in *Stim*^*CA*^ and *Orai*^*CA*^ animals was also quantitatively similar to that seen with *Ras85D*^*V12*^ (43% and 39% for *Stim*^*CA*^ and *Orai*^*CA*^, respectively; Figure 3C,D). Conversely, expression of wildtype *Stim* and *Orai* (*Stim*^*WT*^ and *Orai*^*WT*^, respectively) did not result in significant changes to heart wall thickness compared to controls (Figure 3A-D). *Stim*^*CA*^ expression also resulted in significantly increased heart wall thickness in third-instar larvae (Supplemental Figure S1; https://figshare.com/s/933599115871046a7aee), suggesting that upregulated SOCE can drive excessive cardiomyocyte growth in the developing heart. These results demonstrate that as in mammals, SOCE upregulation results in hypertrophy of the *Drosophila* heart, and that microCT is a valuable tool when combined with genetic approaches for analysis of hypertrophic signaling.

**Figure 3.**
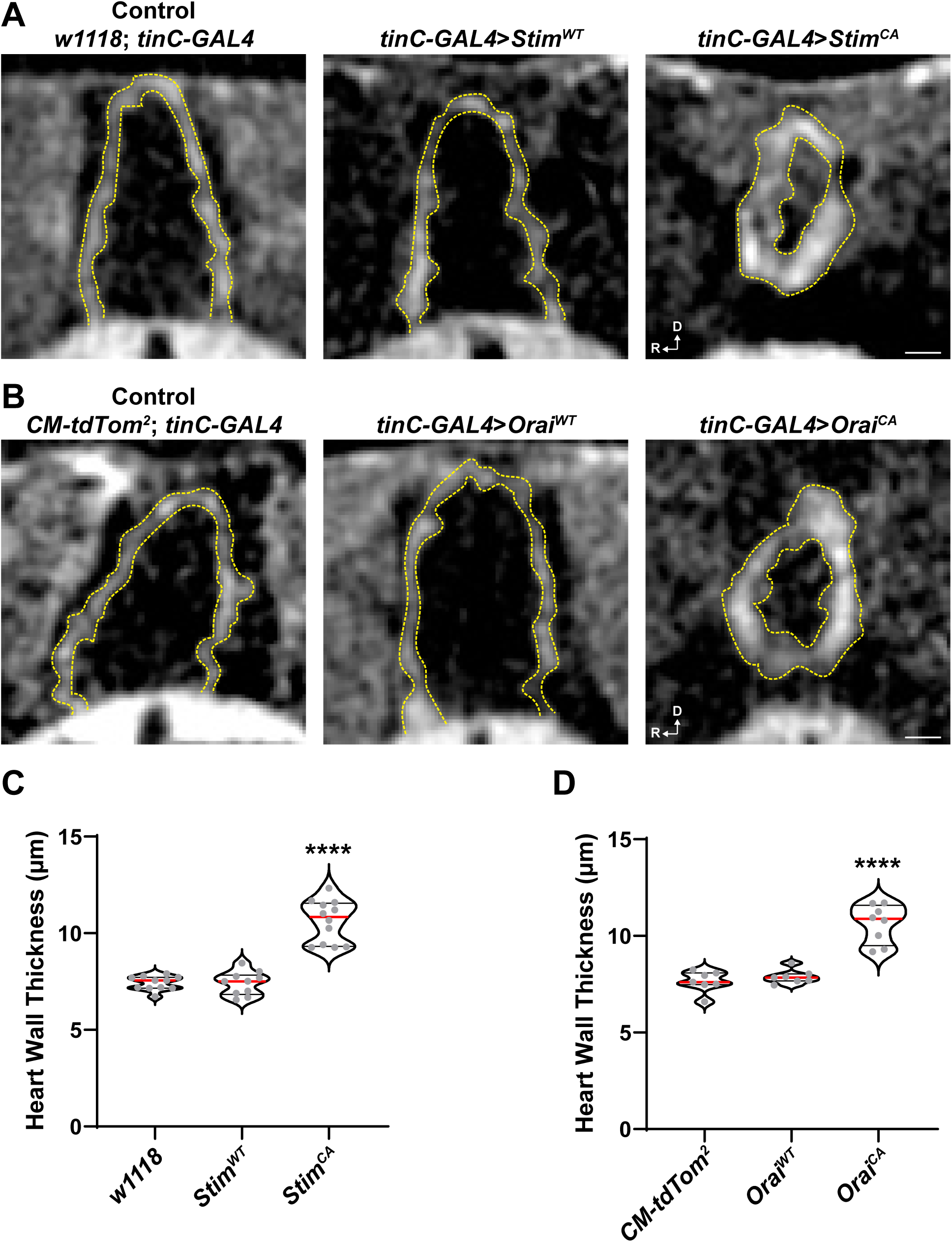
SOCE upregulation causes *Drosophila* cardiac hypertrophy. **A**. Representative images from microCT fast-scans showing heart conical chambers in cross-section from control (*w1118*; *tinC-GAL4*) and *tinC-GAL4* driven *Stim*^*WT*^ and *Stim*^*CA*^-expressing seven day-old females. Dashed yellow lines indicate the inner and outer boundaries of the heart walls. **B**. Representative images from microCT fast-scans showing heart conical chambers in cross-section from control (*CM-tdTom*^*2*^; *tinC-GAL4*) and *tinC-GAL4* driven *Orai*^*WT*^ and *Orai*^*CA*^-expressing seven day-old females (these animals had *CM-tdTom*^*2*^ because the same cross was also used for intravital imaging). Dashed yellow lines indicate inner and outer boundaries of the heart walls. **C**,**D**. Plots of heart wall thickness measurements from fast-scan microCT tomograms of seven day-old females with genotypes shown in panels A and B, respectively. Each circle represents a single animal measurement. Red lines: median; black lines: quartiles. ****p<0.0001 compared to control, one-way ANOVA with Tukey’s multiple comparisons. Scalebars: 25 µm. Anatomical anterior (A) and dorsal (D) are indicated by white arrow coordinates.

Impaired contractility is also a key feature of cardiac hypertrophy. Contractile alterations characteristic of hypertrophy include significantly decreased end-diastolic and end-systolic dimensions due to thickened heart walls, often without changes to fractional shortening. We therefore determined next whether SOCE upregulation impairs contractility of adult *Drosophila* hearts. Heart contractility was analyzed by intravital fluorescence imaging of animals with cardiomyocyte-specific tdTomato (CM-tdTom) expression^26, 28^. Control hearts exhibited a mean end-diastolic dimension (EDD) of 65.17 ± 1.33 μm, mean end-systolic dimension (ESD) of 32.15 ± 0.83 μm, and mean fractional shortening (FS) of 51 ± 0.85% (± SEM; Figure 4A-D). These parameters were not significantly altered by expression of *Stim*^*WT*^ or *Orai*^*WT*^, consistent with the lack of hypertrophy of these hearts by microCT analysis. In striking contrast, EDDs were significantly reduced compared to controls by 26%, 32%, and 15% for *Stim*^*CA*^, *Orai*^*CA*^, and *Ras85D*^*V12*^, respectively (Figure 4A,B). ESDs were also significantly reduced in *Stim*^*CA*^, *Orai*^*CA*^, and *Ras85D*^*V12*^ expressing hearts (Figure 4C), whereas FS was unaltered with *Stim*^*CA*^ or *Orai*^*CA*^ and modestly increased with *Ras85D*^*V12*^ (Figure 4D). Heart rates were similar across control and experimental groups (Figure 4E). These results demonstrate that SOCE upregulation results in contractile dysfunction that is consistent with the hypertrophic phenotype observed in microCT analyses.

**Figure 4.**
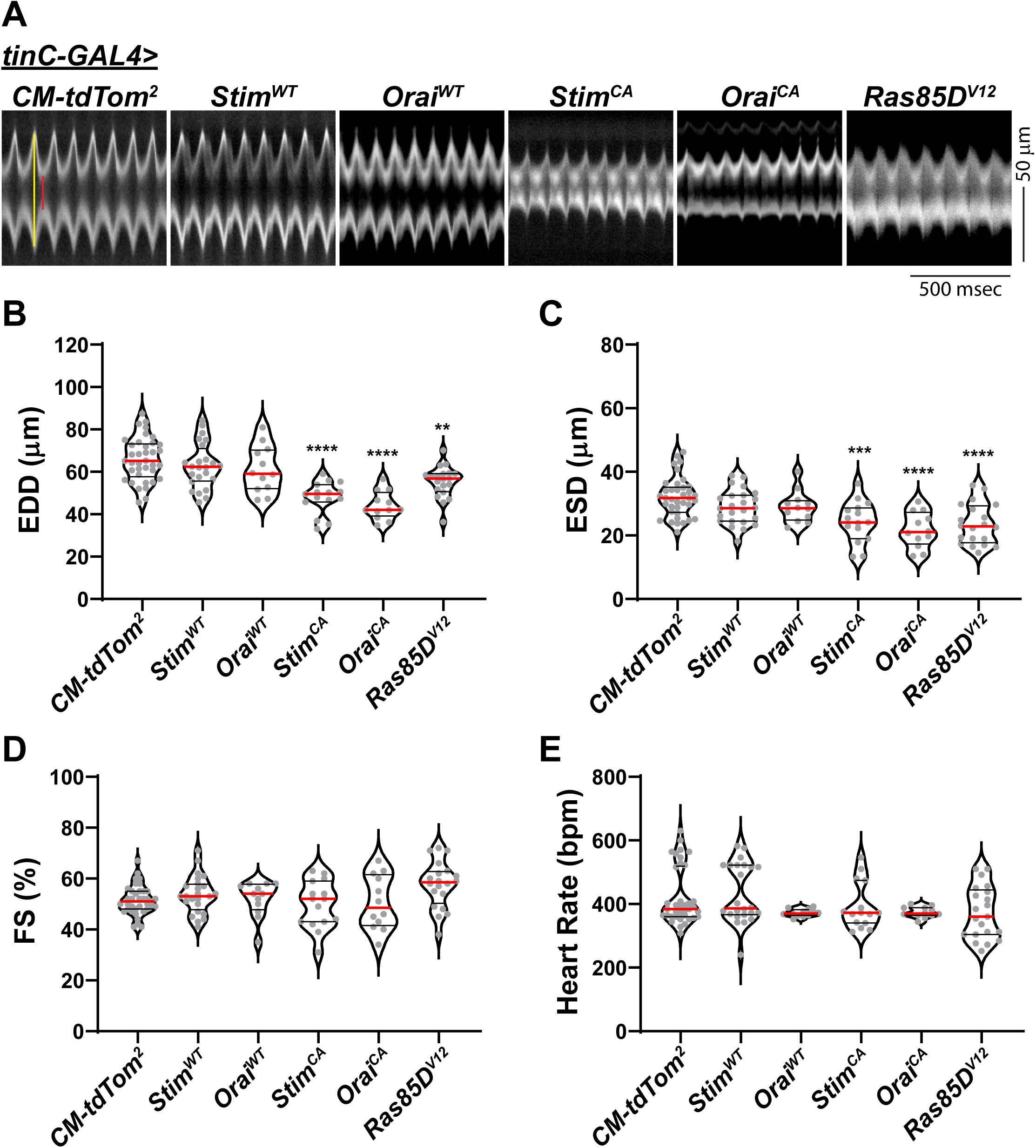
SOCE upregulation impairs heart contractility. A. Heart contractility was analyzed by intravital fluorescence imaging of seven day-old females expressing CM-tdTom. Shown are representative M-modes from animals with *tinC-GAL4* and *CM-tdTom*^*2*^ (control), and *tinC-GAL4* driven *Stim*^*WT*^, *Orai*^*WT*^, *Stim*^*CA*^, *Orai*^*CA*^, and *Ras85D*^*V12*^. The red line in the *CM-tdTom*^*2*^ trace depicts systole and the yellow line depicts diastole. **B-E**. Plots of EDD (**B**), ESD (**C**), FS (**D**), and HR (**E**) calculated from M-modes of the genotypes shown in A. Each circle represents a single animal measurement. Red lines: median; black lines: quartiles. **p<0.01, ***p<0.001, ****p<0.0001 compared to control, one-way ANOVA with Tukey’s multiple comparisons.

### Stim^CA^-mediated cardiac hypertrophy requires Orai channels

As final proof-of-principle for how microCT can be combined with powerful genetic tools to understand heart pathophysiology, we used a genetic suppressor approach to determine whether *Stim*^*CA*^-induced cardiac hypertrophy requires Orai channels. MicroCT analysis again showed significant heart wall thickening in animals with *Stim*^*CA*^ expression alone (Figure 5A,C). Remarkably however, the hypertrophic phenotype was completely suppressed by co-expression of *Orai* RNAi with *Stim*^*CA*^, as heart wall thickness in these animals was similar to controls as well as to animals with *Orai* RNAi alone (Figure 5A,C). This strongly suggests that hypertrophy of *Stim*^*CA*^expressing hearts results from upregulated Ca^2+^ influx through Orai channels. In further support of this conclusion, hypertrophy caused by *Ras85D*^*V12*^ expression was not similarly suppressed by co-expression of *Orai* RNAi (Figure 5B,D). Thus, the ability of *Orai* knockdown to suppress hypertrophy is specific to SOCE upregulation, as opposed to a generalized ability of *Orai* suppression to universally repress hypertrophic growth. Lastly we determined whether the deleterious effects of Stim^CA^ expression on heart contractility similarly depend on Orai channels. In support of this, co-expression of *Stim*^*CA*^ with *Orai* RNAi reversed the reductions in EDD and ESD seen with *Stim*^*CA*^ expression alone (Figure 6A,C-E). Notably, *Orai* RNAi expression alone resulted in significant increases in EDD and ESD compared to controls (Figure 6A,C-E), consistent with dilated cardiomyopathy that results from SOCE suppression^28^. The effect of *Orai* suppression was again specific to *Stim*^*CA*^, as reductions in EDD and ESD caused by *Ras85D*^*V12*^ expression were not similarly reversed by co-expression with *Orai* RNAi (Figure 6B,F-H) and in fact, EDD and ESD were even further reduced compared to *Ras85D*^*V12*^ expression alone. Collectively, these results demonstrate that upregulated SOCE signaling mediated by both Stim and Orai is sufficient to drive cardiac hypertrophy and impaired contractility in *Drosophila*. We further show that the effects of SOCE upregulation are specific and distinct from other mechanisms that drive hypertrophic growth of the heart such as upregulated Ras signaling.

**Figure 5.**
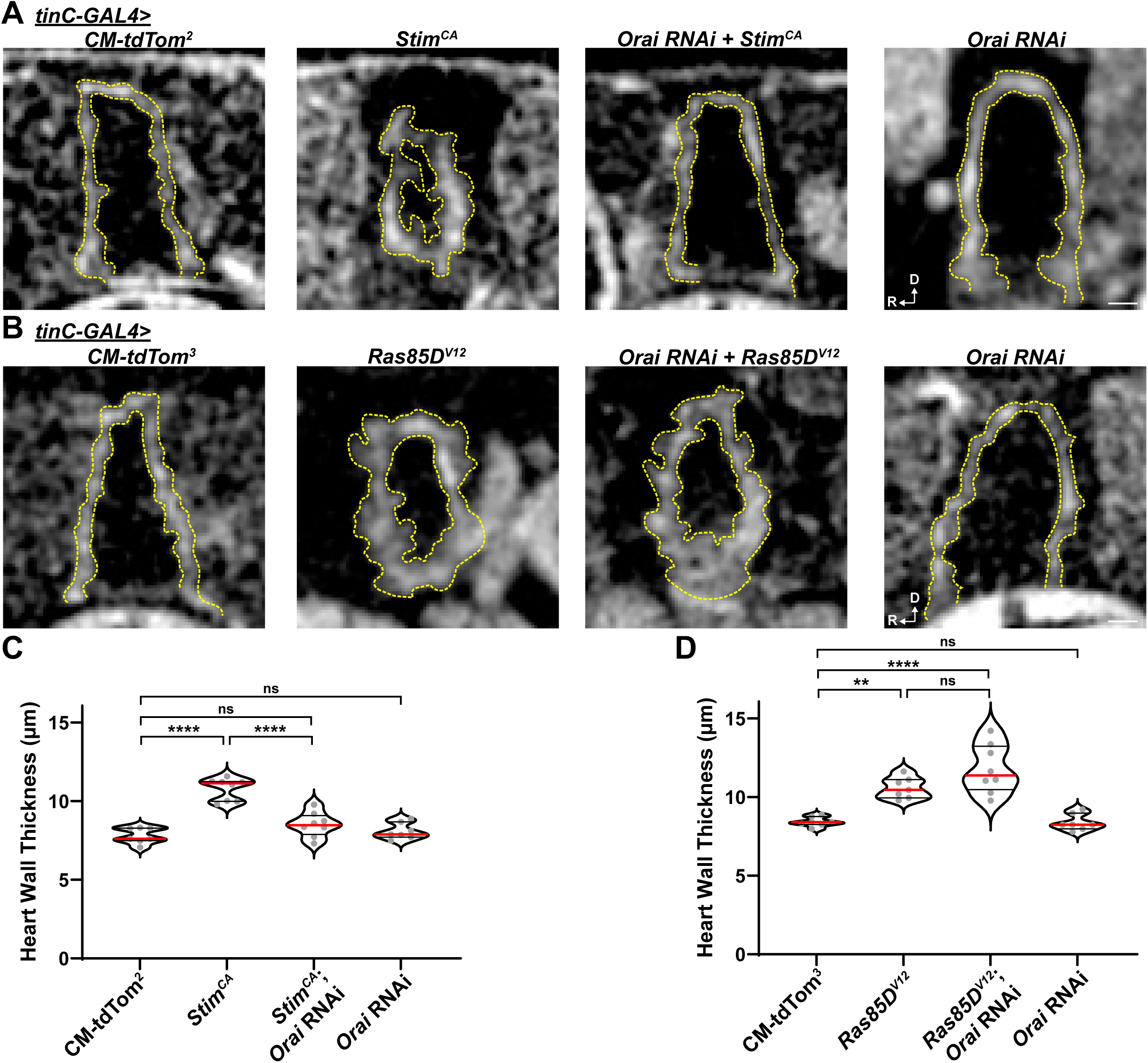
Stim^CA^-driven hypertrophy requires Orai channels. **A**,**B**. Representative microCT fast-scan images showing heart conical chambers in cross-section from control (*CM-tdTom*^*2*^; *tinC-GAL4*) and *tinC-GAL4*-driven *Stim*^*CA*^, *Stim*^*CA*^ + *Orai* RNAi, and *Orai* RNAi expressing seven day-old females (**A**), and control (*CM-tdTom*^*3*^; *tinC-GAL4*) and *tinC-GAL4*-driven *Ras85D*^*v12*^, *Ras85D*^*v12*^ + *Orai* RNAi, and *Orai* RNAi expressing seven day-old females (**B**) (these animals had *CM-tdTom* because the same cross was also used for intravital imaging). Dashed yellow lines indicate inner and outer boundaries of the heart walls. **C**,**D**. Plots of heart wall thickness measurements from fast-scan microCT tomograms of seven day-old females with the genotypes shown in panels A and B, respectively. Each circle represents a single animal measurement. Red lines: median; black lines: quartiles. ns, not significant, **p<0.01, ***p<0.001, ****p<0.0001, one-way ANOVA with Tukey’s multiple comparisons. Scalebars: 25 µm. Anatomical anterior (A) and dorsal (D) are indicated by white arrow coordinates.

**Figure 6.**
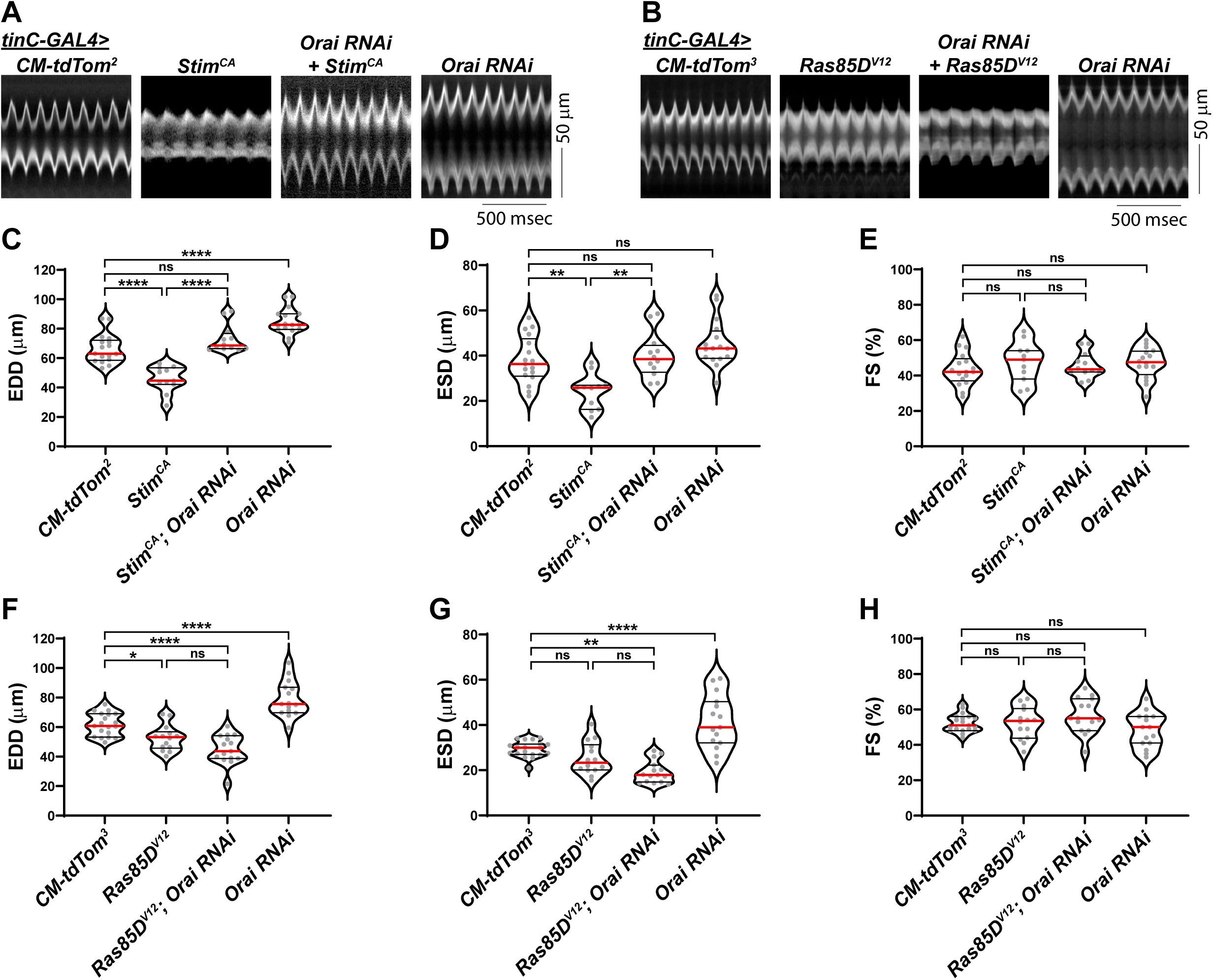
Impaired heart contractility caused by Stim^CA^ expression requires Orai channels. **A**,**B**. Representative M-mode traces from seven day old female flies with *tinC-GAL4* + *CM-tdTom*^*2*^ (control) and *tinC-GAL4* driven *Stim*^*CA*^, *Orai* RNAi + *Stim*^*CA*^, and *Orai* RNAi (**A**), and *tinC-GAL4* + *CM-tdTom3* (control) and *tinC-GAL4* driven *Ras85D*^*V12*^, *Ras85D*^*V12*^ + *Orai* RNAi, and *Orai* RNAi (**B**). **C-E**. Plots of EDD (**C**), ESD (**D**), FS (**E**), calculated from M-mode traces of the genotypes shown in panel A. **F-H**. Plots of EDD (**F**), ESD (**G**), FS (**H**), calculated from M-mode traces of the genotypes shown in panel Each circle represents a single animal measurement. Red lines: median; black lines: quartiles. ns, not significant, *p<0.05, **p<0.01, ***p<0.001, ****p<0.0001, one-way ANOVA with Tukey’s multiple comparisons test.

## DISCUSSION

We have demonstrated that microCT is a powerful, non-destructive imaging platform for analysis of *Drosophila* heart architecture that when combined with genetic tools and functional heart analysis, can be used to decipher complex molecular mechanisms of heart growth and pathological remodeling. Our introduction of microCT imaging to the *Drosophila* heart analysis toolkit has the potential to significantly accelerate our understanding of conserved genetic and signaling mechanisms that regulate heart growth during development and pathological transformations that lead to hypertrophy and heart failure. In contrast to histopathological approaches previously used to analyze *Drosophila* cardiac hypertrophy, we show that microCT yields heart size measurements in flies that are rapidly obtained, highly reproducible, and sensitive enough to detect changes in heart wall thickness indicative of hypertrophy. And importantly, microCT tomograms can be used to assemble complete 3D reconstructions of whole animals, such that the entire heart and other organs can be visualized and measured isometrically in any orientation. Thus, multiple landmarks external to the heart can be used to position consistent measurements, and measurements can always be made from images with the same anatomical orientation. The ability to analyze multiple organs in the same animal with one single dataset also means that microCT offers nearly unmatched potential for studying multi-system diseases that affect the heart as well as other organs^34^.

We validated our microCT approach for detecting *Drosophila* cardiac hypertrophy by analyzing animals that express constitutively-active Ras, an established hypertrophic model based on histopathological analysis^8^. Our microCT heart wall thickness measurements in adult controls were consistent with measurements from histopathological preparations, indicating a normal heart wall thickness of 6-8 μm in the conical chamber. *Ras85D*^*V12*^ expression resulted in a 3-4 μm, or approximately 30-50% increase compared to controls by our microCT measurements, indicating significant hypertrophy of the heart walls. Notably however, prior histopathological measurements showed an approximately 15 μm increase in thickness with *Ras85D*^*V12*^ expression^8^. A possible reason for this apparent discrepancy is that histopathological measurements were made from 8 μm thick heart sections, whereas our microCT measurements were from 2.85 μm optical slices. The increased thickness of histopathological tissue sections may have contributed to the thicker appearance of the heart walls, especially if the hearts were not oriented perpendicular to the plane of section, or if there were heart wall deformations within the section. Thus, we expect that the thinner sections used with microCT result in more accurate measurements of heart wall thickness compared to histopathology.

In addition to heart wall measurements, contractility analysis is also important to defining cardiac hypertrophy phenotypes. Our contractility analyses showed significantly reduced EDDs and ESDs in *Ras85D*^*V12*^, *Stim*^*CA*^, and *Orai*^*CA*^-expressing hearts, consistent with thickened heart walls. It is noteworthy that in microCT images of these hypertrophic hearts, the luminal dimensions of the hearts were significantly reduced compared to controls. This likely reflects the reduced diastolic dimensions measured from contractility analyses. Direct measurement of luminal diameter or area from microCT images could therefore serve as an additional analytical approach to defining hypertrophy. However, we did not attempt these measurements because we have not verified full relaxation of the heart upon fixation. It may be possible to treat animals prior to fixation with agents that stop heart contractility such as Ca^2+^ chelators or Ca^2+^ channel blockers for luminal quantification.

We also demonstrated the utility of microCT for analysis of the larval *Drosophila* heart. This is significant, because the majority of physiological heart growth in *Drosophila* occurs during the larval developmental stages^5, 7, 11^. Thus, microCT may be a valuable tool for delineating physiological mechanisms of developmental heart growth. And because many key signaling mechanisms that drive developmental heart growth are re-activated during pathological cardiac hypertrophy and heart failure, microCT analysis of the larval heart may also be beneficial for uncovering new mechanisms involved in heart failure^35, 36^.

To demonstrate the power of microCT for analysis of signaling pathways that drive cardiac hypertrophy, we used this technique to investigate the role of SOCE signaling in hypertrophy of the *Drosophila* heart. Upregulated SOCE is essential for induction of pathological cardiac hypertrophy^24, 25, 30, 37, 38^. However, we still have much to learn about the precise roles of SOCE signaling in hypertrophic cardiomyocyte growth. Furthermore, a growing number of gain-of-function *STIM1* and *Orai1* mutations have been identified in human patients^39^. Whether these gain-of-function mutations affect human heart physiology is unclear, further highlighting significant gaps in our understanding of SOCE function in the heart. Genetic models of upregulated SOCE are therefore vital to understanding the mechanisms by which SOCE regulates cardiac physiology and disease pathogenesis. Our microCT and contractility data show that genetic SOCE upregulation in cardiomyocytes by *Stim*^*CA*^ and *Orai*^*CA*^ expression results in significant hypertrophy of the *Drosophila* heart, similar to *Ras85D*^*V12*^. The effects of SOCE upregulation on heart function were pathological, as the number of heart-specific *Stim*^*CA*^-expressing animals reaching the pupal stage of development was reduced by approximately 70% compared to controls, and only about 20% of *Stim*^*CA*^ animals reached adulthood (Supplemental Figure S2; https://figshare.com/s/f780cb3d35602163e833).

The similar results observed for both *Stim*^*CA*^ and *Orai*^*CA*^ expression are consistent with a mechanism whereby Ca^2+^ influx through Stim-activated Orai channels is sufficient for induction of cardiac hypertrophy. However, it has been suggested that Stim regulates Orai-independent targets in cardiomyocytes including L-type Ca^2+^ channels and phospholamban^40, 41^. We found that hypertrophy of Stim^CA^-expressing hearts was suppressed by co-expression of *Orai* RNAi, suggesting that Stim^CA^-induced hypertrophy specifically requires Orai channels as opposed to Orai-independent targets in *Drosophila*. In contrast to *Stim*^*CA*^, *Ras85D*^*V12*^-induced hypertrophy was not suppressed by *Orai* RNAi, suggesting that Orai suppression does not have a generalized anti-hypertrophic effect and is specific to hypertrophy caused by Stim activation. This result also suggests that Ras and SOCE function in parallel hypertrophic signaling pathways, as opposed to a linear pathway in which Ras functions upstream of SOCE.

Accumulating evidence suggests that SOCE acts through calcineurin signaling to drive hypertrophic cardiomyocyte growth^23, 24, 30, 37^. However, additional targets of upregulated SOCE in cardiomyocytes have also been reported including CaMKII^23^ and mTORC2/Akt^37^. Thus, consistent with the complex and multi-faceted etiology of pathological cardiac hypertrophy, the role of SOCE in this disease likely involves multiple targets and regulatory processes. Addition of microCT to the *Drosophila* heart analysis toolkit will significantly advance our ability to genetically dissect these complex signaling mechanisms.

## ACKNOWLEDGEMENTS

Reagents obtained from the Bloomington *Drosophila* Stock Center (NIH P40OD018537) and *Drosophila* Genomics Resource Center (NIH 2P40OD010949) were used in this study. JTS was supported by funding from the Collaborative Health Initiative Research Program (CHIRP) of NIH/NHLBI and Uniformed Services University (I 80 VP000003). TAS was supported by the NIH/NHLBI (1 K22 HL137902-01) and an Institutional Development Award (IDeA) from the NIH/NIGMS (2 P20 GM103432).

